# Analysis of DNA methylation in rainbow trout spermatozoa: the strengths and limitations of RRBS

**DOI:** 10.1101/2023.06.20.545730

**Authors:** El Kamouh Marina, Brionne Aurélien, Sayyari Amin, Lallias Delphine, Labbé Catherine, Laurent Audrey

## Abstract

DNA methylation is an important epigenetic mark in fish spermatozoa since it has been shown that some sperm methylome features are transmitted to the offspring. To ensure the transmission of unaltered information to the offspring, the characterization of this mark and its stability in spermatozoa is essential. DNA methylation status can be assessed at the whole genome level with an identification of the methylated and unmethylated cytosines using RRBS (reduced representation bisulfite sequencing). This method allows the sequencing of a subset of the genome expected to be enriched in CpGs. We aim to characterize the data provided by RRBS in rainbow trout spermatozoa, in order to evaluate the suitability of this approach for sperm biotechnologies studies. We observed that RRBS did provide a reduced amount of genomic data, thus allowing the processing of many biological replicates. Although, in our dataset, only a small fraction of the whole genome CpGs was present in all 6 to 12 replicates, the sum of the analyzed CpGs spanned 9 % of the total genomic CpGs. They distributed evenly all over the genome, and all genomic features were represented. RRBS is therefore an effective method to scan the DNA methylation of the along genome in a reduced pattern. However, one should be aware of the choices that are to be made regarding fragment size selection and regarding the options during bioinformatic data processing.

## Introduction

DNA methylation is the most extensively studied epigenetic mark. It involves the covalent binding of a methyl group on a cytosine followed almost exclusively by a guanine in vertebrates. Hence, DNA methylation can be referred to as the methylation of CpG dinucleotides. This epigenetic mark has significant importance in the control of gene expression, X chromosome inactivation and monoallelic gene expression in mammals (Bird 2002). Furthermore, DNA methylation is particularly important in fish spermatozoa. Indeed, it has been demonstrated in zebrafish that some sperm methylome features are transmitted to the offspring and maintained during early embryogenesis (Potok et al. 2013; L. Jiang et al. 2013; Murphy et al. 2018). These features are involved in the control of embryo development. Hence, any alteration of this epigenetic mark in spermatozoa induces the risk of altering the offspring’s development.

To assess DNA methylation at the whole genome level, bulk assay of the cytosines bearing methyl groups include the LUMA (Luminometric Methylation Assay) (Karimi et al. 2006), LC-MS/MS or HPLC (Kurdyukov and Bullock 2016), but these provide a mean methylation level for the whole sample rather than the methylation status of a given cytosine in the genome. Characterization of methylated sites at identified positions in the genome relies mainly on BS-seq methods (DNA bisulfite conversion and sequencing) where unmethylated cytosines are deaminated into uracils which will be converted into thymines after PCR. Since methylated cytosines (both 5-methylcytosine and 5-hydroxymethylcytosine) will not be deaminated, they can be differentiated from thymine-converted unmethylated cytosines after DNA sequencing and mapping to a reference genome. Whole-genome bisulfite sequencing (WGBS) (Frommer et al. 1992), and reduced representation bisulfite sequencing (RRBS) (Alexander Meissner et al. 2005; 2008a; Smith et al. 2009a) are the two genome-wide bisulfite-based methods.

The RRBS method for DNA methylation study differs from WGBS in that it involves endonuclease fractionation of the DNA instead of a random one, and that the number of sequenced fragments is reduced. RRBS was developed for mice genome in order to assess the methylation status of a reduced CpG fraction deemed to be representative of the whole genome (A. Meissner 2005). Using this technique, the sequencing of the whole genome was dispensable. The choice of the MspI restriction enzyme for DNA fractionation further allowed that in mice and humans, the sequenced regions are enriched in CpG sites, namely CpG islands and promoter regions (Smith et al. 2009b; Gu et al. 2010). Hence, RRBS generates, in mammals, a CpG coverage that is higher than the coverage of the sequenced genome fraction. Another advantage of RRBS is that it cuts the genomic DNA at specific restriction sites, and the subset of the fragments resulting from this cut was proposed to be reproducible among the biological replicates (A. Meissner 2005). Last, because RRBS does not involve the sequencing of the whole genome, the amount of data that is produced is reduced. As a consequence, less storage availability and less calculation power for analyses will be needed. This allows, at a lower cost, the analysis of many biological replicates that are very useful when the assessment of sample variability is at stake.

The question of using RRBS for fish genome studies was first settled in zebrafish (Chatterjee et al. 2013) for application to the brain methylome. Despite zebrafish genome being smaller than in humans and mice, the *in silico* testing of RRBS showed that, for the same amount of sequenced genome (about 2.2 % of the whole genome), a lower amount of CpG sites would be recovered in zebrafish compared to humans and mice. This meant that a reduced sequenced fraction would yield a lower fraction of informative CpGs than in humans or mice. This observation raises the question of the suitability of RRBS to really represent enough CpG sites in other fish species.

Rainbow trout is a farmed species with high economic importance for which the transmission of environmental clues to the progeny via DNA methylation in the spermatozoa is an important question. To date, however, only one study assessed sperm DNA methylation with RRBS in this species (Gavery et al. 2018), for the evaluation of genetic and epigenetic variations of sperm from hatchery and natural origin males. While this study provided valuable insights into environmentally-driven genetic and epigenetic modifications in rainbow trout, the efficiency of the RRBS in representing the genomic CpGs sites from a reduced genome fraction was not explored. In the present study, our aim is to characterize the data yielded by RRBS in rainbow trout spermatozoa, in order to evaluate the suitability of this approach on a reduced sequenced fraction of the genome, when the incorporation of many biological replicates (n=12) is at stake.

## Results and Discussion

### Sequencing strategy and mapping efficiency

As shown in Table 1, our RRBS sequencing provided 79 to 182 million raw reads depending on the sperm DNA sample: some males had a much lower number of reads (males 1, 3, 6) than the average while others were over-represented in the sequencing data (males 4 and 8). This variability is due to the difficulty of obtaining a homogeneous representation of the different samples during the sequencing step, after libraries preparation and sample pooling. The males with the lowest read numbers have been resequenced to increase their representation to at least 80 × 10^6^ reads. We surmise that the over representation of some males is due to the hybridization which was randomly more favorable to these samples on the flow cell. Another feature of our data is that among the total raw reads, an average of 53 % had adapters (Table 1). Because our sequencing conditions yielded 100 bases per read, such detection of adapters in more than half the reads means that the smallest DNA fragments (inserts < 100 bp) were abundant in our libraries. The size selection step of the library preparation did not remove enough of the smallest ones, but it cannot be excluded either that the small fragments attached more easily to the flow cell than the bigger ones, so that they became more represented during the sequencing. Since these inserts are < 100 bp and that we carried out a paired-end sequencing, this induced sequencing redundancy between read 1 and read 2 of the same insert. Additionally, the lack of distance between read 1 and read 2 from these too short inserts reduced the chances of the reads to be uniquely mapped to the genome by the Bismark bio-informatic software (Krueger and Andrews 2011). Indeed, we found that an average of 21 % of the trimmed reads were not uniquely mapped, and these were discarded from the analysis.

**Table 1.**
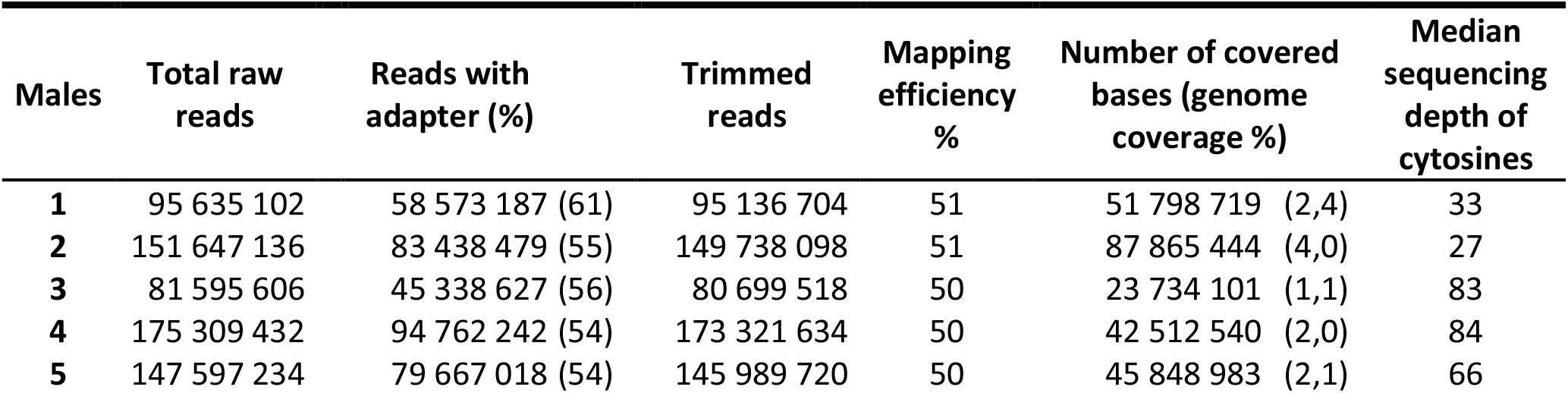
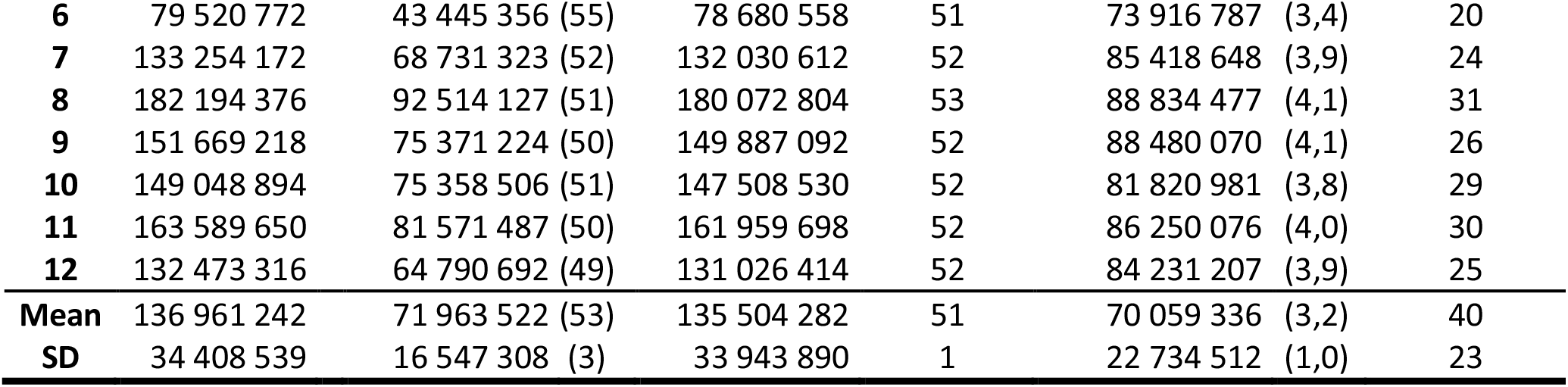
Summary of the sequencing data obtained in our RRBS study. Total raw reads number includes the reads 1 and the reads 2 of each sample (paired end sequencing). Reads with adapter is the number of reads shorter than 100 bp whose adapter was sequenced to reach the 100 bp length of the read. Trimmed reads refer to the number of reads after those shorter than 20 b were removed. Mapping efficiency = (number of reads that mapped only once on the trout genome/trimmed reads number) x 100. Number of covered bases= number of bases that were sequenced and mapped at least once to the reference genome. Genome coverage % = (number of sequenced bases that matched only once with reference genome bases/ trout genome size (2 178 982 971 b)) x 100. Sequencing depth is the number of times a cytosine has been sequenced, mapped to the reference genome and extracted as a cytosine in a CpG context

The mapping efficiency of the reads to the reference genome is another important information when processing RRBS data. Indeed, this parameter reflects the amount of data that will be further analyzed for methylation call and methylation extraction. In all, we observed that only 51 % of the trimmed reads could be confidently and uniquely aligned to the trout reference genome (Table 1). A part of the losses is due to the above-mentioned removal of redundant reads (21 %). Besides, the presence of repetitive sequences in the trout genome, known to account for 57.1 % of the whole genome (Pearse et al. 2019), and the bisulfite conversion of the genome which changed almost all cytosines into thymine bases, thus reducing genome sequence diversity, added up to the alignment difficulties during mapping. Our data revealed that an average of 28 % of reads could not be mapped at all to the reference genome. Thus, our 49 % reads losses during the mapping step consist in non-uniquely mapped reads and reads that were not mapped at all. This low mapping efficiency was also seen in the study that assessed sperm DNA methylation with RRBS in rainbow trout (Gavery et al. 2018), as we calculated that the average mapping efficiency in this work was about 33 % of the total reads. RRBS data are not the only sequencing data affected by this phenomenon as a recent study using whole genome bisulfite sequencing (Brionne et al. 2023) also showed a reduced mapping efficiency with this trout genome (approximately 60 % of the reads).Our low mapping efficiency was also observed after RRBS study in the spermatozoa of some mammalian species with an average mapping efficiency of only 19 % in cattle (Z. Jiang et al. 2018), 55 % in human (Ingerslev et al. 2018), whereas it was up to 70 % in rats (Suvorov et al. 2018).

The last parameter describing the sequencing yield with RRBS is the number of bases from the reference genome that were sequenced in our experiment. When considering all the reads that were uniquely mapped to the reference genome, a mean genome coverage of 3.2 % was obtained (Table 1), but this coverage was highly variable among the males. It is noteworthy that the males who had the lowest raw reads are not necessarily the ones with the lowest genome coverage (see male 6). With our set of data, male 3 represented only 1.1 % of the reference genome. We infer that for this specific sample, many reads represented the same genomic regions, and this is further emphasized by the high CpGs sequencing depth (times the same CpG site is sequenced) for this sample (n=83) despite its low read number. Such low genome coverage in one sample may become a problem when comparisons with the other males are sought, and this male may have to be discarded from further analysis.

### Information yielded by RRBS about the genomic CpGs

Among the 2 178 982 971 bases of the rainbow trout reference genome, there are 35 336 285 CpG dinucleotides (Omyk_1.0 genome version). At the molecular level, this number of CpG is the same on the forward and reverse strand, and we will refer to them as CpG sites (in which we include the cytosine in CpG context on the forward strand, and its corresponding one on the reverse strand). But during the building of the libraries, both forward and reversed strands can be represented (that can be called plus and minus strands when referring to the reference genome), and this can result in the sequencing of the same CpG site twice (once on the forward strand, once on the reverse strand). As a consequence, the treatment of the data can be carried on in two ways. First, we can consider that the methylation status of the cytosines at a given CpG site is the same on both strands. This is the case in most situations where a biological process is studied, since DNA methylation occurs symmetrically and it can be repaired by DNA methyltransferase in the case of hemi methylated DNA. Thus, the redundancy between the two strands must be removed during the data treatment, and the resulting unique information is one out of the 35 336 285 CpG sites. Second, we can be interested in the methylation status of the cytosines on each strand of a given CpG site. This is the case for example when some biotechnologies are applied to the spermatozoa. If no repair mechanism has been operating, a CpG site can then be altered on the forward strand but not on the reverse one. Therefore, each CpG site is characterized by the methylation status of two cytosines (one from the forward and once from the reverse strand). As a consequence, the information is kept whole and is one out of 70 672 570 cytosines (twice the number of genomic CpG sites). This seemingly fastidious choice is important in the case of RRBS whose sequencing yield is much lower than by WGBS. In our set of data, we calculated that an average of 28 % of the analyzed cytosines belonged to the reverse strand of a forward cytosine. This can lead to either 28 % redundancy that is to be removed from the data (first choice) or 28 % of sites for which a possible hemimethylation can be explored (second choice).

### The question of enrichment in CpG sites after RRBS

Because RRBS reduces the sequencing extent, as confirmed above, we sought which proportion of the CpG sites of the trout genome were present in our data. Our purpose was also to investigate whether RRBS in rainbow trout spermatozoa would capture DNA fragments enriched in CpG sites. As shown in Table 2, 2 to 6.9 % of the trout genomic CpG sites were represented depending on the male. As this value is meant per CpG site, the cytosines found on both the forward and reverse strand have been calculated as one single information. However, when we performed the same calculation over the 2 cytosines from each CpG site of the genome, we obtained a lower representation (from 1.2 to 5.2 % depending on the males, Table 2). This is due to the fact that, as mentioned above, only 28 % of the sequenced cytosines belonged to both DNA strands for a given CpG site. When the percentage of genomic CpG sites covered in our analysis (Table 2) was compared to the percentage of genome coverage (Table 1), in order to assess the enrichment in CpG sites among the sequenced fraction, we observed a very homogeneous enrichment of 1.7-1.8 X in CpG sites in all males. The enrichment fold was only 1.2 X when counting the 2 cytosines from each CpG site. Hence, in the later situation, RRBS does not provide an enrichment in analyzed cytosines, but it is still providing some information on a small fraction of the genomic CpGs.

**Table 2.** Number of CpGs obtained by RRBS compared to the total CpGs in the rainbow trout genome. % total genomic CpG sites = (number of identified CpG sites after RRBS / number of CpG sites in the reference genome (35 336 285)) x 100. Enrichment in CpG sites/genome = % total genomic CpG sites / % genome coverage. The % genome coverage values for each sample are available in the Table 1. Number of identified cytosines: includes all the cytosines in CpG sites whatever their sequencing depth. The % total genomic cytosines in a CpG context refers to (number of identified cytosines in CpG sites / number of cytosines in a CpG context in the reference genome (70 672 570)) x 100

| Males | Number of identified CpG sites | % total genomic CpG sites | Enrichment in CpG sites/genome | Number of identified cytosines | % total genomic cytosines in a CpG context |
| --- | --- | --- | --- | --- | --- |
| 1 | 1 532 633 | 4,3 | 1,8 | 2 031 111 | 2,9 |
| 2 | 2 406 116 | 6,8 | 1,7 | 3 667 465 | 5,2 |
| 3 | 710 888 | 2,0 | 1,8 | 813 505 | 1,2 |
| 4 | 1 244 325 | 3,5 | 1,8 | 1 514 634 | 2,1 |
| 5 | 1 323 326 | 3,7 | 1,8 | 1 611 036 | 2,3 |
| 6 | 2 044 500 | 5,8 | 1,7 | 2 898 053 | 4,1 |
| 7 | 2 369 361 | 6,7 | 1,7 | 3 559 536 | 5,0 |
| 8 | 2 444 130 | 6,9 | 1,7 | 3 679 864 | 5,2 |
| 9 | 2 422 373 | 6,9 | 1,7 | 3 665 894 | 5,2 |
| 10 | 2 263 435 | 6,4 | 1,7 | 3 292 897 | 4,7 |
| 11 | 2 364 475 | 6,7 | 1,7 | 3 536 955 | 5,0 |
| 12 | 2 300 074 | 6,5 | 1,7 | 3 405 609 | 4,8 |
| Mean | 1 952 136 | 5,5 | 1,7 | 2 806 380 | 4 |
| SD | 591 835 | 1,7 | 0,1 | 1 027 725 | 1 |

These enrichment values are the result of a sequencing experiment on biological samples, where the exact size of the sequenced DNA fragments (inserts) cannot be precisely controlled. We compared these data to a theoretical in *silico* digestion at MspI sites (CCGG) analysis that we performed on rainbow trout genome (Table 3). Our objective was also to explore whether inserts size would affect this enrichment ratio. We observed that theoretical fragments of increasing size (40-220 b to 150-1400 b) led to decreasing theoretical enrichment ratios (Table 3), from 2.5 X with the smallest fragments to 1.3 X with the biggest ones. This means that when selecting bigger fragments for sequencing, the enrichment ratio is reduced, and this would hint that most CpGs are in the smallest fragments fraction of the genome. This is further confirmed by the distribution of the CpGs according to the size of the fragments (Fig. 1), as we found that the majority of the CpGs were in the small fragments fractions. It is nevertheless difficult to compare the predicted *in silico* enrichment values to the 1.7 X enrichment ratio that we obtained with our sequenced data. Indeed, although the sequencing company proposed a fragment size selection of 150-1 400 b, which should have yielded an enrichment of 1.3 X, we know that much smaller fragments were sequenced (< 100 b, see the discussion above). This illustrates the gap between the theoretical fragments size eventually proposed by the sequencing company and the size of the fragments which were really sequenced.

**Table 3.** *In silico* analysis of the rainbow trout genome upon RRBS treatment. A Msp1 digestion of the reference genome of rainbow trout was performed *in silico*, and the fragments were *in silico* selected according to different size ranges (Selected fragments size, in bases). (1) comparison with the zebrafish *in silico* analysis (Chatterjee et al. 2013) and (2) the mouse one (calculated from (Smith et al. 2009b)). RR genome size (Mb) is the number of bases obtained *in silico* when selecting a given fragments size range. % genome coverage = RR genome size/genome size x 100. % CpG = number of CpG sites/ number of CpG sites in the reference genome x 100. CpG sites enrichment ratio = % CpG / % genome coverage

| Species | Rainbow trout |  |  | Zebrafish (1) | Mouse |
| --- | --- | --- | --- | --- | --- |
| Selected fragments size (b) | 40 - 220 | 40 - 400 | 150 -1 400 | 40 - 220 | 40 - 22 |
| Genome size (Gb) | 2.18 | 2.18 | 2.18 | 1.41 | 2.8 |
| RR genome size (Mb) | 56 | 142 | 676 | 31 | 38 |
| % genome coverage | 2.6 | 6.5 | 31.1 | 2.2 | 1.4 |
| Number of CpG sites | 2 241 808 | 4 436 904 | 13 820 867 | 1 430 390 | 1 506 7 |
| % CpG | 6.3 | 12.6 | 39.11 | 5.3 | 7.0 |
| CpG sites enrichment ratio | 2.5 | 1.9 | 1.3 | 2.4 | 5.0 |

**Fig. 1.**
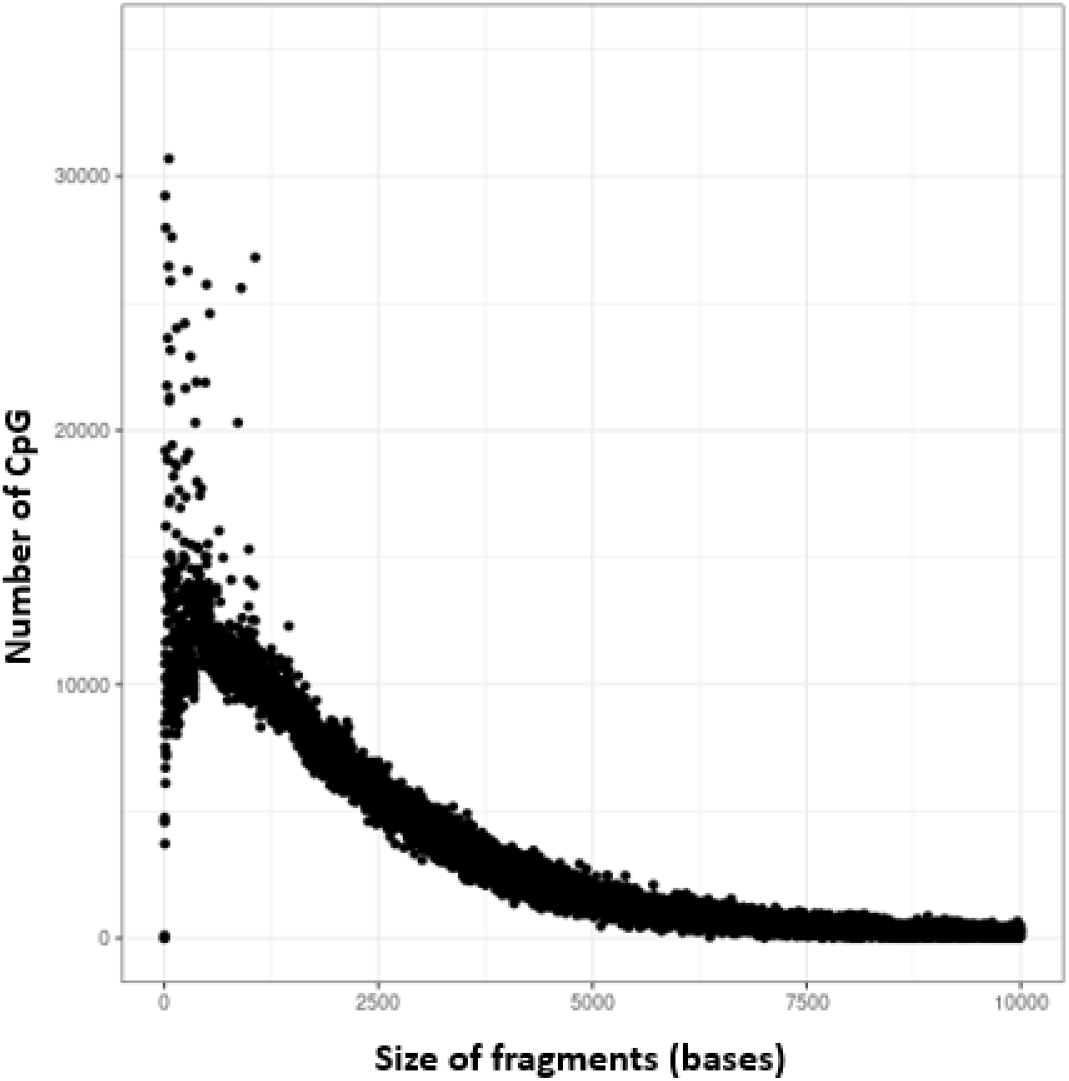
Distribution of the CpG sites number according to the DNA fragment size. Each dot represents the total number of CpG sites in all the fragments of a given size. For the sake of representation, 3 dots corresponding to 3 extreme CpG number values have been hidden: 52 133 CpGs for 27 b fragments, 41 640 CpGs for 42 b fragments and 37 587 CpGs for 52 b fragments. The number of CpGs and the size of the fragments were obtained from our *in silico* digestion of the rainbow trout reference genome (Omyk_1.0 genome version)

In zebrafish, it has been shown *in silico* that RRBS induced a CpG sites enrichment of 2.4 X when the selected fragments were between 40-220 bp (Chatterjee et al. 2013), a value that is very similar to the one that we obtained *in silico* with the rainbow trout genome (Table 3). These two calculations show that RRBS did not provide the same enrichment as it did for mammals. Indeed, with the same selection of size fragments, the enrichment was 5 X in mice (Smith et al. 2009b) and 5.9 X in human (Chatterjee et al. 2013). Such low CpGs enrichment in fish can be explained by the fact that fish genomes are less heterogeneous regarding CpG density than in mammals (Cai et al. 2021). This would reduce the chance of obtaining CpG dense regions in the fish genome fragments, leading to a restricted genomic CpG representation.

In all, our thorough analysis of CpG enrichment after RRBS in rainbow trout genome, both experimentally and *in silico*, emphasizes the compromise that is to be made when using this method. The sequencing of the smallest fragments should theoretically enrich the fractions with CpG sites, but these small fragments will poorly map to the reference genome, and will be sequenced twice (read 1 and read 2) but to no avail due to the later on removal of the redundant information. Bigger fragments will better map to the reference genome, their read 1 and read 2 will not overlap, but their CpG density will be lower. This compromise is important to clarify before carrying on the step of libraries preparation and fragment size selection.

### Representation of RRBS CpGs in biological replicates

Since RRBS consists in DNA digestion with a restriction enzyme, the fragments that will be selected and sequenced are more likely supposed to represent the same genomic regions among biological replicates, as previously discussed by (Alexander Meissner et al. 2008b) for mammalian genomes. This should yield CpG sites that will be more similar among biological replicates than would random fragmentation, such as the one obtained after sonication for example. Hence, we explored in our study whether the represented genomic CpG sites were repeatable among the biological replicates. Since this issue is important when assessing the impact of reproductive biotechnologies on CpG methylation in fish, we determined the genomic CpG sites representation at the cytosine level (the forward and reverse ones in a CpG site), thereby using all 70 672 570 cytosines of the CpG sites of the reference genome (plus and minus strands). We found that an overall 9.1 % of the cytosines in genomic CpG sites were represented in our data set (6 422 403 cytosines at CpG sites), when all the cytosines were combined irrespective of the male origin of the data. However, at the male level, an average of 4 % only were represented per sample (Table 2). This can be explained by the fact that, contrarily to the expectations in other species or studies (Alexander Meissner et al. 2008b), a high number of fragments sequenced in some samples were not sequenced in the others. We then investigated further to which extent the same cytosines were repeatably observed or not between the different males. As shown in Fig. 2, the highest proportion of cytosines among the 6 422 403 analyzed ones was present in one male only. Interestingly, about the same proportion of cytosines were common between 3 to 9 males, although their number was reduced (8.1 to 9.3 % of the 6 422 403 analyzed cytosines). The numbers dropped when selecting the common cytosines to 10 to 12 males, the lowest value for 12 males being due to the low cytosine yield in male 3 (Table 2). This shows that both distinct and common information was obtained between biological replicates. Nevertheless, the low proportion of common cytosines between males (< 10 % of the analyzed ones) indicates that the fragments that were sequenced was very different between males, despite the non-random fractionation of the genome using MspI restriction enzyme. A small part of this phenomenon could be due to the genetic polymorphisms at CpG sites among these biological replicates, resulting in fragments that would be different between males. However, this polymorphism cannot account for the whole extend of the divergent cytosines between biological replicates. Another explanation could again be due to the too many small fragments that were randomly selected. We infer that a size selection of bigger fragments would have yielded a smaller number of fragments of the required size, and thus a better overlapping of these fragments between males upon sequencing.

**Fig. 2.**
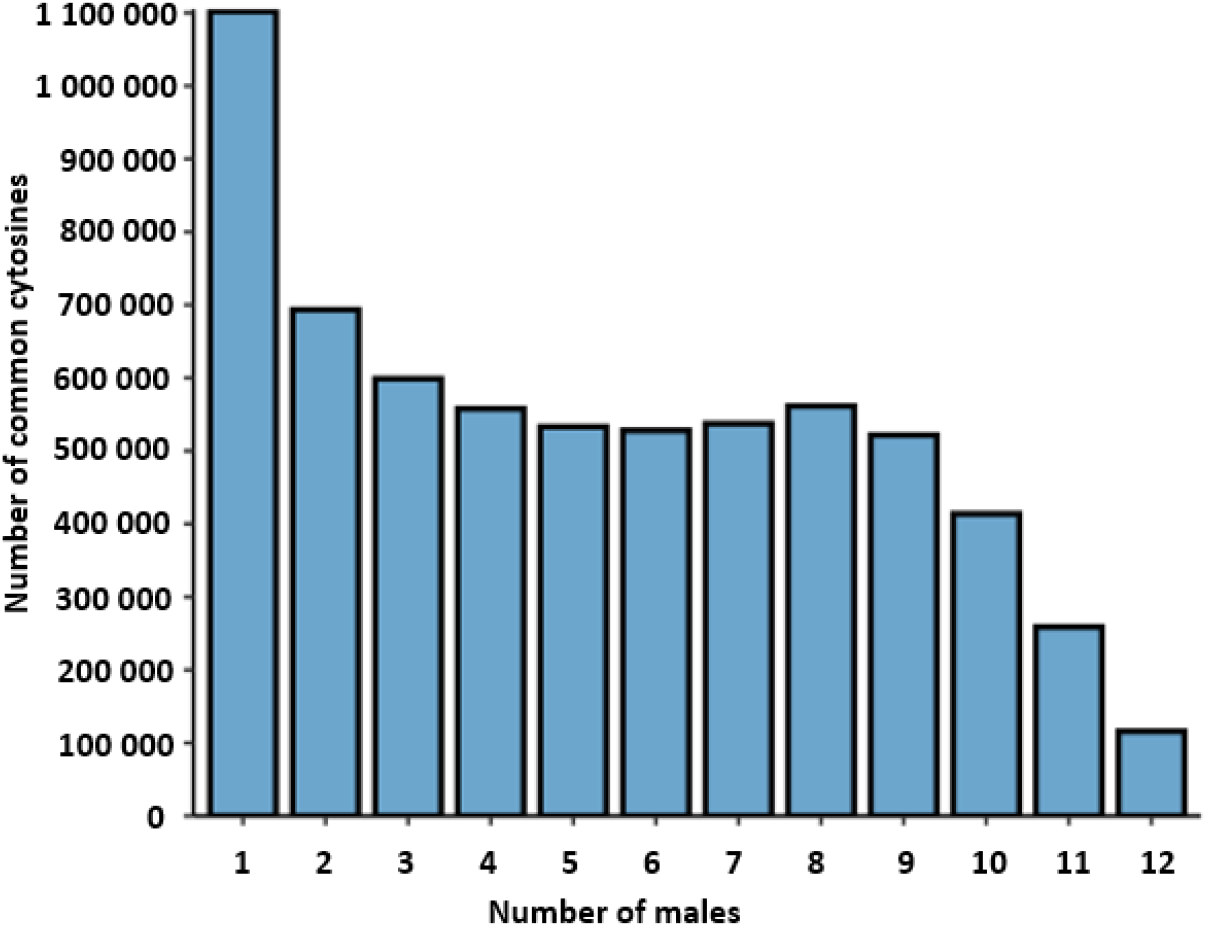
Distribution of the number of cytosines in a CpG context that are common between biological replicates. Each sequenced cytosine (in CpG context) was checked for the number of male samples where it was found (from 1 to 12). Bars show the distribution of the number of common cytosines according to the number of males. Number of common cytosines in 1 male refers to the cytosines that were unique

In all, these results showed that increasing the number of replicates in RRBS analysis ensured an acceptable representation of the genomic cytosines in CpG sites. However, it also showed that any study of the variability between biological replicates would rely on a very reduced number of cytosines that would be common between males, at least in our conditions. This also stresses once more the importance of the fragment size selection step and strategy.

### Distribution of the data over the reference genome

All our analyses concurred to show that RRBS provides a reduced representation of the genome with only a moderate enrichment in CpG sites. Hence, we sought to determine the distribution pattern of the sequenced CpG sites over the whole genome, in order to ensure that the genome is homogeneously represented with our data. It is particularly important when assessing the effect of biotechnologies, because all regions possibly affected, including the random ones in non-functional genome regions, are of interest. As shown in the example Fig. 3, our sequenced cytosines (in CpG sites) were homogeneously distributed all over the genomes. In addition, we did not identify any genomic regions bigger than 2.9 × 10^6^ bases that were devoid of these sequenced cytosines. Our results also demonstrated that the represented genomic CpGs per sample were distributed homogeneously across all the chromosomes of the genome. This means that despite the restricted number of CpGs in the RRBS data set, they give some information all over the genome.

**Fig. 3.**
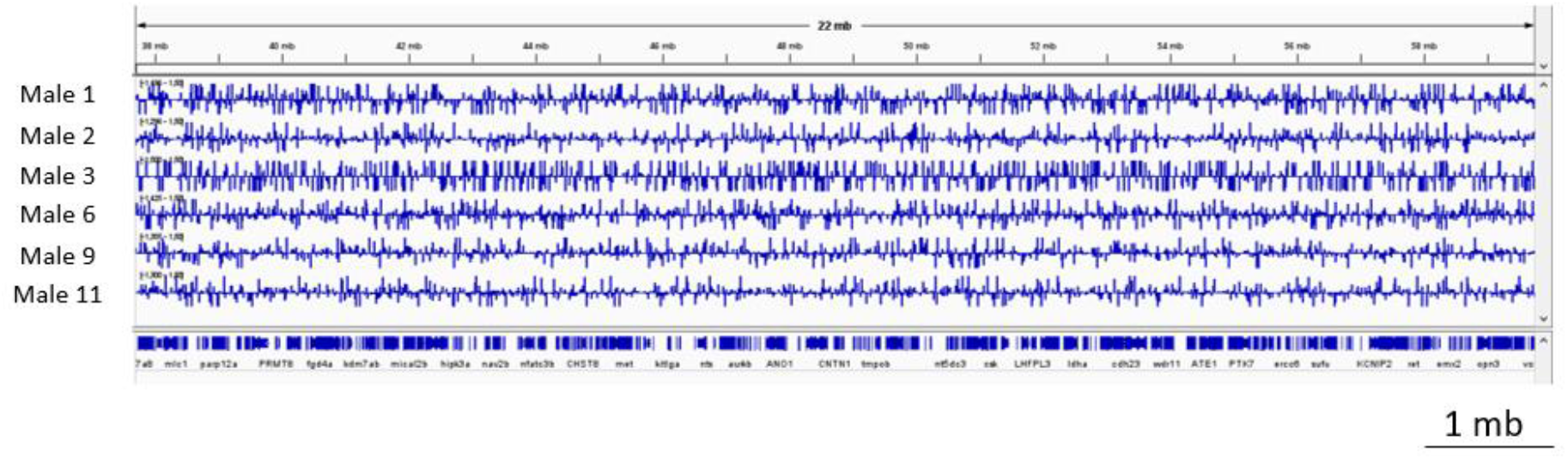
Distribution of the sequenced cytosines in CpG sites over the reference genome. The six males represented here have been randomly chosen. The displayed region is representative of the pattern observed over the whole genome. The chart was obtained with IGV_2.8.13 and shows the distribution on the chromosome 1 from the position 37 682 451 to 59 737 137 (22.10^6^ b) of the rainbow trout reference genome “Omyk_1.0 genome version”. Each line corresponds to a different male, where each sequenced cytosine in a CpG site is represented by a blue bar. Long and short bars represent methylated and unmethylated cytosines respectively. The bars position above and below the horizontal line represent cytosines on the forward and reverse strands respectively. The bottom line represents the genes associated to this genomic fraction. Scale bar: 10^6^ bases

To investigate further the genome representation of our RRBS data, we assessed to what extent the sequenced genomic CpG sites belonged to the different genomic features that are promoter and downstream regions, 5’ and 3’UTR regions, introns and exons or intergenic regions (Fig. 4). We observed that the CpG genomic sites obtained in our RRBS data were found in all genomic features, most of them being found in gene introns and intergenic regions. Besides, this distribution was repeatable between males, as observed from the low variability of the data. When we compared the distribution of the analyzed CpGs that are annotated into these genomic features in each sample to the one of WGBS (Brionne et al. 2023), we found that the distribution of the analyzed CpGs resulting from RRBS in each sample was similar to the one obtained with WGBS data (Fig. 4), the later representing 92 % of the reference genome CpGs. This suggests that although the different total number of CpGs analyzed per male between the 2 methods, the distribution of the CpGs into genomic features was not affected by the genome representation extent. It is noteworthy however that CpG distribution in exons tended to be higher in our RRBS data than in WGBS ones. On the contrary, the CpGs distribution in promoter was lower in RRBS than in WGBS. Although no statistics can be drawn from this comparison, this discrepancy may deserve further attention regarding a possible method-related bias in the representation.

**Fig. 4.**
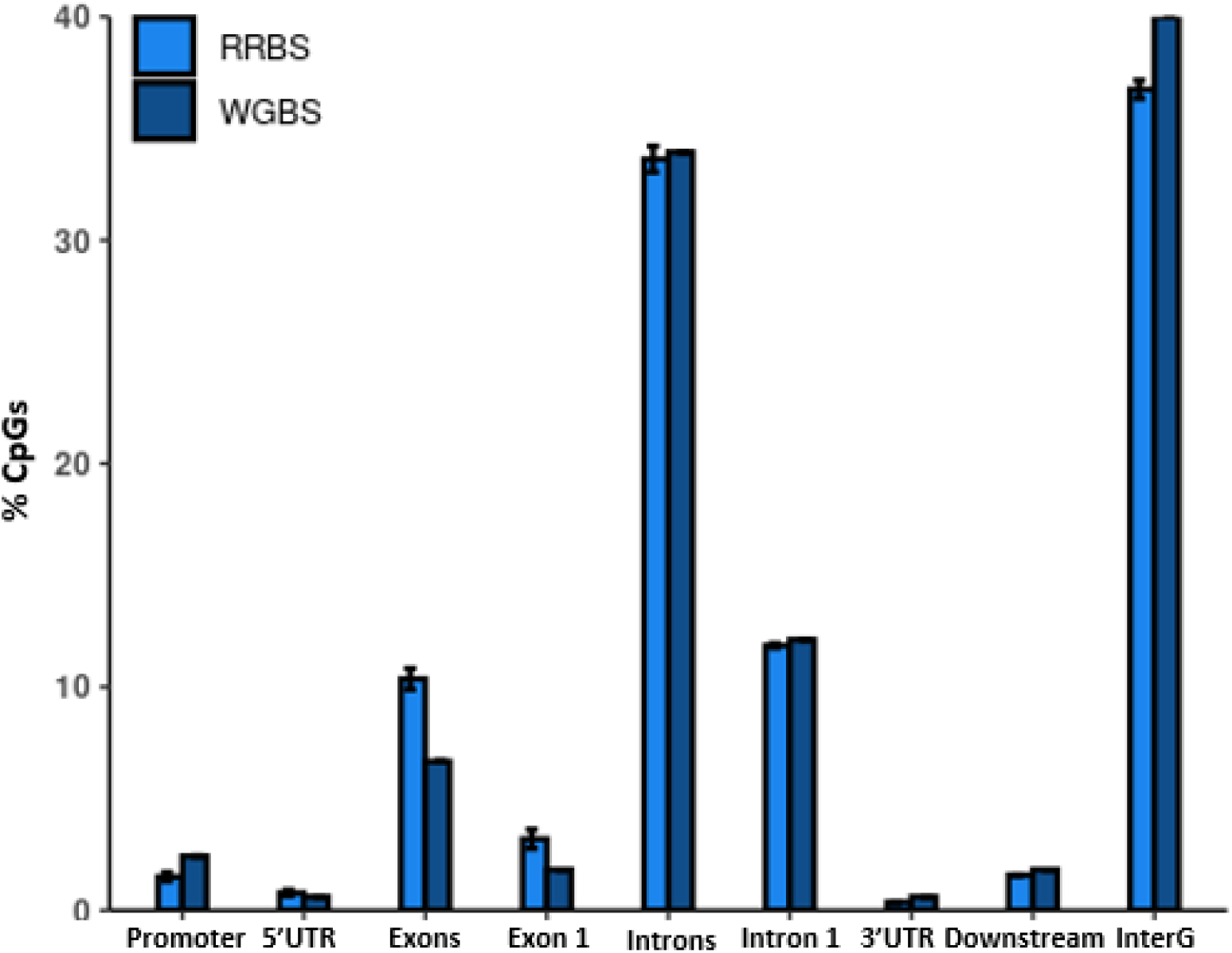
Distribution of the CpGs sites obtained by RRBS according to their genomic features, and comparison with WGBS published data. For RRBS, bars represent the mean value of the 12 biological replicates. The values are calculated as a percentage of all the cytosines present in at least one of the chosen genomic features. Error bars: standard deviation. For WGBS, data were calculated from (Brionne et al. 2023). The chosen genomic features were: Promoter: -1kb up to the transcription starting site (TSS); 5’UTR: untranslated region upstream of the first translated codon; Exons: all exons; Exon 1: first exon of all genes; introns: all introns; 3’UTR: untranslated region downstream of the STOP codon; Downstream: transcription end site (TES) +1kb; InterG: intergenic regions

Based on the homogeneous distribution of CpGs along the genome and the representation of different genomic features by these CpGs, RRBS represented the whole genome in a reduced but representative pattern.

### The risk of random male comparisons

In the present work, all males belonged to the same genetically diverse line and they were all reared in the same conditions from fertilization to spermiation. Additionally, all sperm samples were processed similarly. As a consequence, all sperm samples should bear the same general DNA methylation profile, although some individual variability would still be present (genetic polymorphism and individual physiology). We took the set of data from these biological replicates to build two sham groups of 6 fish, each fish having been randomly allotted to a given group. Each group thus contained individual samples with slightly variable information, but such variability should be smoothened when processing the data as a group. As a consequence, no difference between groups should arise from group comparison. However, when we used DSS for the 2 groups comparison, we obtained as many as 1 593 differentially methylated cytosines (DMCs). This striking observation means that when one wants to compare sperm samples for different treatments for example, a background of about 1600 DMCs may have no ground for biological significance. This illustrates the risk of overestimating the number of DMCs when comparing treatment to control conditions. One specificity of our data analysis is that we wanted to keep the raw DNA methylation status at each CpG position. As a consequence, we disabled the smoothing option of “dmltest” function in DSS package, and the methylation values were then either close to 0 % or close to 100 %. These yes-or-no values may have been favorable to such spurious DMC detection.

A deeper analysis of the significant DMCs additionally showed that these were detected from heterogeneous replicate numbers between groups, the outermost values being as little as one male in each group, or one male in one group and 6 males in the other group. This means that for a given cytosine position, some DMCs were identified as significant when only one male in one group differed from 1 to 6 males in the other group. We were able to make this striking observation because the RRBS data were small enough to be hand-made and controlled with regards to the number of males involved in each significant DMC. When we applied a threshold so that each DMC should be found in at least 5 males in each group, the number of these spurious DMCs dropped to 22, a more acceptable (because negligible) value. This illustrates why it is to be recommended that the number of biological replicates at each CpG position should be carefully set before the analysis.

This sham analysis emphasizes the need to carefully adapt the bioinformatic parameters to the dataset and to the experimental purpose, and that one should not go blindly into data interpretation on the basis of DMCs that may not bear any biological significance. One great advantage of studying the effect of biotechnologies on sperm is that in most cases, the same sperm sample can be split between control and treatment conditions. This allows the use of the paired comparison option in DSS, thus strengthening the biological significance of the identified DMCs and preventing the bias identified in the present study.

## Conclusion

Our study showed that DNA methylation analysis of rainbow trout sperm by RRBS provided some information on the methylation status of about 10 % of the genomic CpG sites, provided that enough biological replicates were analyzed (n=12 in our case). These sites were evenly distributed all over the genome, and they were present in all genomic features. It can thus be considered that the reduced representation provided a widespread, although restricted, picture of genomic CpGs in rainbow trout. We also emphasized that the method provided only a limited enrichment in CpG site compared to the genome coverage, as already shown in zebrafish, and contrarily to human and mice. Besides, our data contained very little overlapping of CpG sites between biological replicates (< 10 % of the analyzed sites), making the study of inter-individual variability very sparse. Last, our data showed the difficult compromise that is to be made between the sequencing of small fragments enriched in CpGs but difficult to map, and that of bigger fragments with less CpGs but easier to analyze. The choice of the statistical parameters set for data analysis should also be carefully chosen to avoid any bias in data interpretation. In all, RRBS does provide a reduced representation of the genomic CpGs in rainbow trout, making possible the analysis of many biological replicates, but with little access to inter-individual variability, at least in our experimental and analytical conditions.

## Materials and methods

### Ethics statement

Rainbow trout males were kept in the experimental fish facilities ISC LPGP of INRAE (Agreement number D-35-238-6) with full approval for experimental fish rearing in strict accordance with French and European Union Directive 2010/63/EU for animal experimentation. The Institutional Animal Care and Use Ethical Committee in Rennes LPGP (Fish Physiology and Genomics Department) specifically approved this study (n° T-2020-37-CL). All fish were handled for gamete collection in strict accordance with the guidelines of the Institutional Animal Care and Use Ethical Committee in Rennes LPGP (Fish Physiology and Genomics Department). Catherine Labbé is accredited by the French Veterinary Authority for fish experimentation (n° 005239). The animal study is reported in accordance with ARRIVE guidelines (https://arriveguidelines.org) for animal research.

### Sperm collection and DNA extraction

Two years old mature males from the SYNTHETIC INRAE strain (average weight 1.2 kg) reared at the experimental fish farm of INRAE-PEIMA were transferred to INRAE-ISC LPGP fish facility. After 2 weeks in recirculated water tanks (12 °C, winter photoperiod), the sperm from 12 males was collected by gentle striping and stored individually on ice. After sperm concentration determination by flow cytometry, 4 × 10^6^ spermatozoa per sample were snap frozen and stored at -80 °C until use. Thawed spermatozoa were digested overnight at 42°C in 800 μL TNES buffer and proteinase k (P6556 ; Sigma-Aldrich) prior to DNA extraction according to (Depincé et al. 2020). DNA concentration was determined by the Qubit™dsDNA BR Assay kit (REF Q32853, Invitrogen by Thermo Fisher Scientific) and 500 ng was sent for RRBS library preparation and sequencing at the Integragen company (Evry, France).

### RRBS library preparation, sequencing and data processing

DNA digestion with MspI, library preparation and bisulfite conversion using Diagenode Premium RRBS kit were performed in July 2021 by the Integragen company, and the samples were sequenced on Illumina NovaSeq™ 6000, with S2 patterned flow cells. The number of sequenced clusters that we ordered to the company was calculated from the following: the size of the rainbow trout genome is 2.18 Gb and the genomic representation that we sought by RRBS was 5 % of the genome, yielding to 0.109 Gb to be sequenced. We knew from Brionne et al (2023) that the mapping efficiency would be about 50 % with the trout genome. As a consequence, we had to double the number of bases to be sequenced (0.218 Gb). Because we chose a 20-25X sequencing depth, we calculated that we should sequence 5 Gb per sample. Since the sequenced reads are 100 b long, the 5 Gb corresponds to 50 × 10^6^ raw reads. This led us to order 25.10^6^ paired-end raw reads. Because of some difficulties with the first sequencing round (too many reads with inserts), some samples have been resequenced after some changes in fragment size selection. This yielded an average of 68.5 million paired end raw reads, 137 million total raw reads (Table1). The sequencing quality of the raw data was checked with FastQC program (version: FastQC_v0.11.7) (Andrews, n.d.). Raw reads were trimmed using Trim Galore to remove adaptors sequences, the bases added during RRBS library preparation, and the bases with a phred score < 30 (https://github.com/FelixKrueger/TrimGalore version: TrimGalore-0.6.5).

In order to determine if there is a risk of overestimation of DNA methylation due to a low bisulfite conversion (< 98 %), we calculated the bisulfite conversion rate and found a mean value of 99.5 %. Using bowtie2 aligner (version: bowtie2-2.3.5.1) available in Bismark bio-informatic software (Krueger and Andrews 2011), trimmed reads were aligned to the reference genome (Omyk_1.0 genome version). Methylation status of every cytosine in a CpG context was extracted using “bismark_methylation_extractor”. These steps are available in the workflow which agrees to FAIR principles and is accessible online (https://forgemia.inra.fr/lpgp/methylome). BigWig files were obtained with bedGraphToBigWig (Kent et al. 2010) and methylation values were visualized with Integrative Genome Viewer (Robinson et al. 2011).

For the identification of differentially methylated cytosines (DMCs), the “2 group comparison” analysis from DSS R package (Feng, Conneely, and Wu 2014; Wu et al. 2015) was used. Each group had 6 males chosen randomly and all the cytosines in CpG context of each sample were used (regardless of their sequencing depth). In addition, we used the option “smoothing=FALSE” for the dmlTest function performed by DSS, leading to the absence of any smoothing of DNA methylation status between adjacent CpGs. This allows that, in a context of studying technological impact, any sporadic alteration remains visible.

For the in silico analysis of RR genome, we performed an in silico digestion with MspI on the Omyk_1.0 genome version using DECIPHER v2.0 R package (Wright 2023). We used “DigestDNA” available in DECIPHER v.2.0 package in order to cut the DNA on the forward strand at the CCGG site. We also specified that the cut takes place after the first cytosine of this site. These options were applied to mimic the digestion performed with MspI enzyme. “DigestDNA” also showed the fragments positions. “Biostrings: Efficient manipulation of biological strings” R package (Pagès et al. 2023) was used to identify the fragments size and CpG content.

## Acknowledgment

The authors acknowledge the skillful involvement of the staff from the ISC LPGP and UE INRAE PEIMA for animal rearing and care. The authors wish to warmly thank Vincent Coustham for his valuable comments on the interpretation of our RRBS data which triggered the motivation to write this paper.

## Statements and Declarations

### Funding

MEK is recipient of a PhD fellowship ARED from region Bretagne and INRAE PHASE (2020-2023). AS obtained the financial support from the internal funding scheme at the Norwegian University of Life Sciences (project number 1211130114), which financed AS international stay at INRAE, Fish Physiology and Genomics, UR 1037, Rennes, France. This work was funded by the French CRB Anim project ANR-11-INBS-0003 and the European FEAMP BIOGERM measure 47 (Innovation Aquaculture).

## Competing Interests

The authors declare no competing interests.

## Author contribution

MEK organized and carried out the study, analyzed and interpreted the data, and drafted the manuscript. AB taught and supervised the bioinformatic work of MEK, and contributed to most bio-informatic analyses. AS contributed to the wet lab work. DL contributed to the conception of the project and provided her knowledge of the trout genome for data interpretation. AL and CL conceived, designed and supervised the study. CL supervised the writing of the manuscript and AL provided her knowledge on genomic DNA methylation in trout. All authors approved the final version of the manuscript.

## Data availability

The data for this study has been deposited in the European Nucleotide Archive (ENA) at EMBL-EBI under accession number PRJEB60579 (https://www.ebi.ac.uk/ena/browser/view/PRJEB60579).

## Notes

### Competing Interest Statement

The authors have declared no competing interest.

